# *Xanthomonas protegens* sp. nov., a rice seed-associated probiotic and taxonomic outlier species of *Xanthomonas sontii*

**DOI:** 10.1101/2024.04.23.590712

**Authors:** Rekha Rana, Anushika Sharma, Vishnu Narayanan Madhavan, Ramesh V. Sonti, Hitendra K. Patel, Prabhu B. Patil

## Abstract

Rice seed microbiota play a vital role throughout their growth and developmental stages. *Xanthomonas sontii* is known as an abundant, core-vertically transmitted rice seed endophyte with probiotic properties. Studies reveal a less explored world of non-pathogenic *Xanthomonas* (NPX) species like *X. sontii* and *X. indica* that are associated with healthy rice seeds. The present study reports the isolation, biochemical characterization, and in-depth taxonogenomic evidence for a third NPX species, *X. protegens*, from rice seeds, a taxonomic outlier of *X. sontii*. It also highlights the importance of using multiple taxonogenomic indices to identify taxonomic outlier species. Similar to the other two NPX species, the new member species is also non-pathogenic and provides *in vivo* protection to rice plants from *Xanthomonas oryzae* pv. *oryzae* upon leaf clip inoculation. The pangenome investigation suggested a set of unique genes to the novel species that might be important in adaptation to rice plants and differentiate it from other NPX species. The study will allow the design of markers for the identification of these NPX members, targeted culturomics of a particular NPX species and in the discovery of other novel NPX species from healthy rice seed microbiome for further exploitation in rice health.

## Introduction

*Xanthomonas* group of bacteria are primarily known as serious phytopathogens (Ryan *et al*., 2011). However, with the advancement of genomics and culturomics, xanthomonads are increasingly recognised as non-pathogenic members of healthy plant microbiome (Cesbron *et al*., 2015, Bansal *et al*., 2021, Rana *et al*., 2022). Rice is the staple food crop for more than half of the world’s population. In the last century, only one species of *Xanthomonas* was known in rice as the causal agent of bacterial leaf blight (BLB), a serious disease of rice (NIÑOLLIU *et al*., 2006). However, in the last decade, three new non-pathogenic *Xanthomonas* (NPX) species have been reported from the rice microbiome. *X. maliensis* was the first NPX species reported to be associated with healthy rice plant, which was isolated from leaves (Triplett *et al*., 2015). Further, two more NPX species, *X. sontii* and *X. indica*, were reported from healthy rice seeds (Bansal *et al*., 2021, Rana *et al*., 2022). There have been many studies using metagenomics and culturomics focussing on the microbiome associated with healthy rice seeds (Cottyn *et al*., 2001, Midha *et al*., 2016, Eyre *et al*., 2019). These studies revealed the presence of strains from the genus *Xanthomonas* that have probiotic properties, such as siderophore and phosphate utilization, seedling development, and biocontrol (Raj *et al*., 2019, Ahumada *et al*., 2022, Zhang *et al*., 2022). This suggests that seeds of rice and other plants harbour several NPX species that form a community and play an important role as the seed microbes are the first inhabitants and are associated with the plant throughout the lifecycle of the host.

Interestingly, most of the species from the genus *Xanthomonas* currently fall in the clade II group of Xanthomonads, which primarily comprises serious pathogens (Koebnik *et al*., 2021, Mafakheri *et al*., 2022). However, many of the recently reported novel species belong to the clade I group of xanthomonads (Bansal *et al*., 2021, Mafakheri *et al*., 2022, Rana *et al*., 2022, Chuang *et al*., 2023). Most of these species follow a non-pathogenic lifestyle and lack major virulence factors such as T3SS and T6SS and their effectors. Clade I of xanthomonads is comparatively less explored as it has fewer economically important pathogens. With the reduced cost of genomics, more and more new studies are being conducted in this group. These novel species might explain the evolutionary trajectory of the genus *Xanthomonas* and might also benefit host plants by helping with nutrient acquisition and fighting pathogens (Fang *et al*., 2015, Rana *et al*., 2023). One of these NPX species, *X. sontii*, was recently reported as an abundant, core, vertically transmitted endophytic, keystone species of rice seeds with numerous probiotic properties (Zhang *et al*., 2022, Rana & Patil, 2023). In another study, early inoculation of *X. sontii* led to the recruitment of several taxa of beneficial bacteria and allowed their networking, reinforcing the importance of NPXs (Wang *et al*., 2023). Hence, there is an urgent need to understand the community of NPX in general and rice in particular. Such studies will allow us to understand the evolutionary success of each of the species and also develop markers in identification and genes unique to each of these species for further molecular genetic studies. In the present study, we report the isolation and characterization of a novel NPX member species from healthy rice seeds. Like other NPX species of rice, i.e., *X. sontii* and *X. indica*, the new member species is also able to protect rice from bacterial leaf blight pathogen upon leaf clip inoculation (Rana *et al*., 2023). Interestingly, the new NPX species is also the closest taxonomic relative of *X. sontii* and forms a complex with *X. indica*. The new species will allow systematic studies to understand the importance of the NPX community in rice seeds and the success of *X. sontii* as a keystone species. Further, a genome-based phylogenetic tree and pangenome analysis will help us confirm the borderline species.

## Material and methods

### Bacterial isolation

Strains were isolated from healthy rice seeds obtained from Gujarat in 2021. The isolation protocol was the same as in the literature (Midha *et al*., 2016). Briefly, the rice seeds were crushed in sterile pestle mortar after removing the husks using sterile forceps. Crushed seeds were dissolved in autoclaved 0.85% saline solution and incubated for 2 hours at 28 °C at 180 rpm. The solution was serially diluted up to 10^-5^ dilution and plated on four different media, i.e., Nutrient Agar (NA), Peptone Sucrose Agar (PSA), Tryptone Sucrose Agar (TSA), and Glucose Yeast extract Calcium carbonate Agar (GYCA). The plates were incubated at 28 °C for 3-5 days. Yellow mucoid colonies were selected and streaked on NA to obtain pure cultures. Strains were stored in 15% glycerol at −80 °C.

### 16S rRNA gene sequencing and phylogeny

The partial 16S rRNA gene sequences were PCR amplified using universal primers 27F (5′-AGAGTTTGATCCTGGCTCAG-3′) and 149R (5′-GGTTACCTTGTTACGACTT-3′) using ready-to load 5X FIREPoL® master mix (Solis BioDyne). The PCR conditions were: initial denaturation for 5 minutes at 95 °C, 30 cycles of denaturation (40 s at 95 °C), annealing (45 s at 55 °C) and extension (1 min 30 s at 72 °C) and a final extension of 10 min at 72 °C. The amplified PCR product was sent to Eurofins Genomics India Pvt. Ltd. (Bengaluru) for sanger sequencing using 27F primer. The partial 16S rRNA gene sequences were identified using the EzBiocloud webserver (www.ezbiocloud.net) and submitted to the NCBI GenBank database (www.ncbi.nlm.nih.gov/).

For phylogenetic analysis,16S rRNA gene sequences were extracted from draft genomes using barrnap v0.9 (https://github.com/tseemann/barrnap), except *X. maliensis* M97 (KF992843) and *X. arboricola* LMG747 (Y10757), which were fetched from LPSN (www.bacterio.net). The sequences were aligned with partial deletion of gaps using the MUSCLE program, and a Neighbor-Joining phylogenetic tree was constructed in MEGA-X v10.2.6 using the jukes-cantor model of substitution with 1000 bootstrap replicates (Edgar, 2004, Kumar *et al*., 2018). The phylogeny was edited and colour-coded in iTOL v6 (Letunic & Bork, 2021).

### Transmission Electron Microscopy

PPL118^T^ was grown on NA and subsequently inoculated in Nutrient Broth (NB) at 28 °C at 180 rpm overnight. The cells were pelleted down by centrifugation at 2000 rpm for 10 minutes. The pellet was washed twice in Phosphate Buffer Saline (PBS) and resuspended in it. The bacterial suspension was positioned on a carbon-coated grid for 15 minutes. The grid was negatively stained with 2% phosphotungstic acid for 30 sec, dried and observed under a JEM 2100 transmission electron microscope (JEOL, Tokyo, Japan) operating at 200kV.

### Biochemical characterization

The biochemical characterization of PPL118^T^ was performed using the BIOLOG GEM III microplate system (Biolog Inc, USA), as per protocol A provided by the manufacturer. The BIOLOG plates were inoculated with the culture incubated at 28 °C. Three different plates were incubated on three different days, and three readings were taken for each plate. The plates were read using a MicroStation 2 reader at 24 h, 48 h, and 72 h, and the output was interpreted using MicroLog 3/5 2.0.1 software. Only +/- values were considered valid. Readings taken at 48 h were more consistent in comparison to 24 h and 72 h.

### DNA isolation and whole genome sequencing

Genomic DNA was isolated from overnight grown cultures of PPL118^T^, PPL117, PPL124, PPL125, and PPL126 in NB using Quick-DNA™ Fungal/Bacterial Miniprep Kit (Zymo Research, USA). Nanodrop 1000 (Thermo Fisher Scientific) was used for quantitative and qualitative assessment. The DNA was shipped off to MedGenome (Hyderabad) for whole genome sequencing using their Illumina NovaSeq platform facility. The paired-end raw reads were assessed for their quality and presence of adapter sequences using FastQC v0.11.9 (https://qubeshub.org/resources/fastqc). The adapter sequences and reads below the Phred score 30 were discarded using Trim-galore master v0.6.7 (Krueger, 2015) with Cutadpat v1.18 (Martin, 2011). Trimmed paired-end reads were *de-novo* assembled using SPAdes v3.15.5 (Prjibelski *et al*., 2020). The assembled genome parameters, such as genome coverage, GC content (%), and N50 values, were determined using Quast v5.0.2 (Gurevich *et al*., 2013). Genome completeness and contamination were calculated using CheckM v1.2.2 (Parks *et al*., 2015). The assembled genomes were submitted to the NCBI GenBank database and were annotated using NCBI’s Prokaryotic Genome Annotation Pipeline (www.ncbi.nlm.nih.gov/genome/annotation_prok/).

### Taxono-phylogenetic analysis

The Average Nucleotide Values (ANI) were estimated using oANI (Lee *et al*., 2016) applied with USEARCH v11.0.667 (Edgar, 2010) and were further confirmed using ANIb (ANI blast) calculated using JSpeciesWS (Richter *et al*., 2016). The digital DNA-DNA hybridization (dDDH) values were calculated using formula 2 of Genome to Genome Distance Calculator v3.0 (Meier-Kolthoff *et al*., 2021). For phylogenetic analysis, genomes of representatives of the genus *Xanthomonas* were fetched from the NCBI GenBank database. Prokka v1.14.6 was used for genome annotation (Seemann, 2014), and Roary v3.13.0 was used for obtaining core gene alignment at protein identity 90% with default parameters (Page *et al*., 2015). The phylogenetic tree was constructed using PhyML v3.3.20211231 with 1000 bootstrap replicates and Generalised Time Reversible (GTR) model of nucleotide substitution (Guindon *et al*., 2010).

### Pan-genome analysis and virulence factors

Anvi’o platform v7.1 was used for the pan-genome analysis of PPL118^T^, *X. sontii* PPL1^T^, *X. sacchari* CFBP4641^T^, and *X. indica* PPL560^T^ (Eren *et al*., 2015). DIAMOND (Buchfink *et al*., 2015) and MCL algorithm (Van Dongen & Abreu-Goodger, 2012) were used to estimate gene similarity and predict gene clusters, respectively. AntiSMASH v7.0.1 was used to predict the Biosynthetic gene clusters (BCGs) in the genomes (Blin *et al*., 2023). The Lipopolysaccharide gene cluster was extracted from genomes using a homology search of *etfA* and *metB* genes, which are situated at the ends of the gene cluster (Patil & Sonti, 2004). The gene sequences *etfA* and *metB* were taken as queries in tBLASTn v2.12.0+ against genomes taken as subjects. The visual maps of the gene clusters were constructed using Clinker v0.0.23 (Gilchrist & Chooi, 2021). The presence of secretion systems and related factors in PPL118^T^ was predicted using Macsyfinder v2.1.3 with the “TXSScan” model (Abby & Rocha, 2017).

### Plant inoculation assays

To test the pathogenicity against rice plants, PPL118^T^ was inoculated in rice leaves. Milli-Q and *X. oryzae* BXO43 were taken as negative and positive control, respectively. Briefly, the cultures were grown overnight in Peptone Sucrose (PS) media, pelleted down, washed with Milli-Q water and resuspended in Milli-Q, O.D._600nm_ adjusted to 1. 60 to 80 days old rice leaves of Taichung Native 1 (TN1) cultivar were clip inoculated with the strains. Further, to check whether PPL118^T^ provides protection against bacterial blight disease development due to *X. oryzae*, co-inoculation was also performed. For this, BXO43 and PPL118^T^ resuspended in Milli-Q at O.D._600nm_ 1 were mixed in the ratio 1:1 to make up the final volume of 1 ml. The mixture was clip-inoculated in 60 to 80 days old TN 1 rice leaves. The disease lesion was measured 14 days post-inoculation.

## Results and discussion

### Strain isolation, identification, morphology, and 16S rRNA gene phylogeny

Five isolates, PPL117, PPL118^T^, PPL124, PPL125, and PPL126, were isolated from healthy rice seeds obtained from Gujarat. The isolates form pale yellow-coloured colonies after 24 h to 48 h grown on NA/PSA at 28 °C. PPL118^T^ cells were rod-shaped with single polar flagella (**Supplementary Fig. 1**). The partial 16S rRNA gene sequences showed more than 99% similarity to *Xanthomonas sontii* PPL1^T^ for all the strains using the EzBiocloud database. The second closest hit was *X. indica* type strain, PPL560^T^. The partial 16S rRNA gene sequences are submitted to the NCBI. In the 16S rRNA gene-based phylogeny, the novel strains, PPL118^T^, PPL117, PPl24, PPL125, and PPL126, are forming a monophyletic clade in a subclade consisting of *X. sontii* strains. This subclade is further clustered in clade I of Xanthomonads with another non-pathogenic species reported from rice, *X. indica* strains and sugarcane pathogen *X. sacchari* strains (**Fig. 1**). Based on 16S rRNA gene identity and phylogeny, the isolates were identified as diverse strains of *X. sontii*.

**Figure 1.**
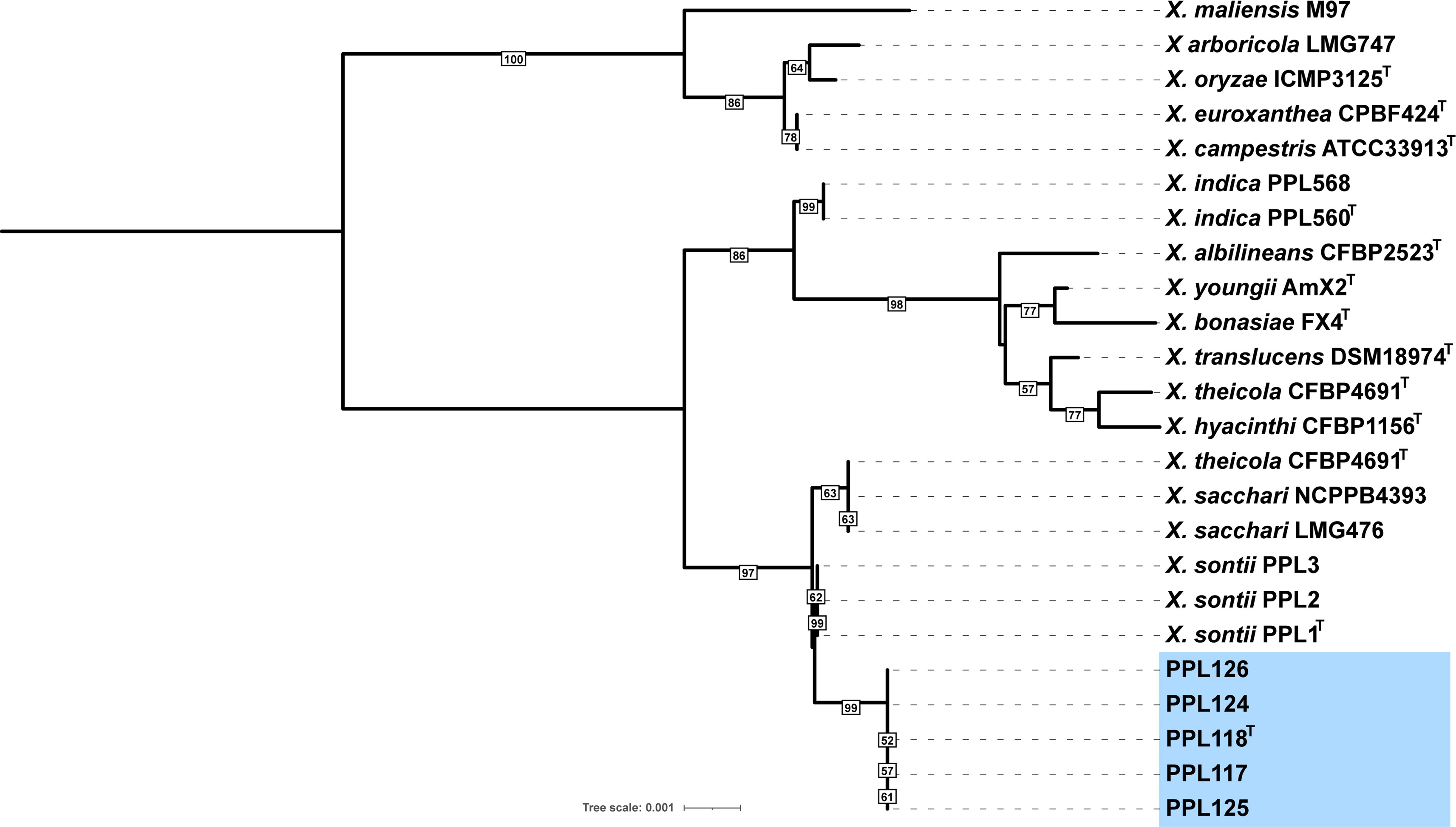
A mid-rooted Neighbor Joining phylogeny constructed using 16S rRNA encoding gene sequences of the novel isolates with representatives of other important species of the genus *Xanthomonas*. The novel isolates are highlighted in light blue colour. The bootstrap values are mentioned on the tree branches in black bordered boxes (values below 50% are filtered out). The tree scale represents the number of nucleotide substitutions per site.

### Phenotypic characterization of PPL118^T^

Biochemical tests using 95 substrates were performed for carbon source utilization, enzymatic activity, and acid production. PPL118^T^ is able to utilize Dextrin, D-Maltose, D-Trehalose, D-Cellobiose, Gentiobiose, Sucrose, D-Turanose, α-D-lactose, D-melibiose, β-methyl-D-glucoside, Gelatin, D-fructose, D-galactose, α-D-glucose, D-mannose, L-alanine, L-aspartic acid, L-glutamic acid, L-lactic acid, citric acid, Bromo-succinic acid, Propionic acid, Tween 40, and Acetic acid. PPL118^T^ was resistant to Rifampicin SV, Lincomycin, Vancomycin, Tetrazolium violet and Tetrazolium blue. The strains showed growth at pH6, 1% NaCl, and 1% sodium acetate. The complete biochemical profile of PPL118^T^ is given in **Supplementary Fig. 2**. The biochemical profile of PPL118^T^ was also compared with its closest species, *Xanthomonas sontii*, which has the same biochemical properties except for L-fucose utilization (**Table 1**). The novel isolate PPL118^T^ shares major phenotype characteristics with its closest relative *X. sontii*, indicating it is similar yet different from this important species.

**Table 1:** Phenotypic characterization of PPL118^T^ in comparison to *X. sontii* PPL1^T*^ (*data taken from literature). Symbols represent +/- for source utilization/for lack of metabolism, and v for variable readings.

| S. No. | BIOLOG tests | PPL118<sup>T</sup> | *X. sontii* PPL1<sup>T</sup> |
| --- | --- | --- | --- |
| 1 | Dextrin | + | + |
| 2 | D-maltose | + | + |
| 3 | D-Trehalose | + | + |
| 4 | D-Cellobiose | + | + |
| 5 | Gentiobiose | + | + |
| 6 | Sucrose | + | + |
| 7 | D-turanose | + | + |
| 8 | D-raffinose | v | - |
| 9 | α-D-lactose | + | + |
| 10 | D-melibiose | + | + |
| 11 | β-methyl-D-glucoside | + | + |
| 12 | Gelatin | + | + |
| 13 | D-fructose | + | + |
| 14 | D-galactose | + | + |
| 15 | L-fucose | - | + |
| 16 | L-rhamnose | - | - |
| 17 | D-sorbitol | - | - |
| 18 | D-mannitol | - | - |
| 19 | D-arabitol | - | - |
| 20 | L-alanine | + | + |
| 21 | L-aspartic acid | + | + |
| **22** | L-glutamic acid | + | + |
| **23** | Formic acid | v | v |
| **24** | Quinic acid | + | + |
| **25** | Citric acid | + | + |
| **26** | Bromo-succinic acid | + | + |
| **27** | Propionic acid | + | + |
| **28** | Acetic acid | + | + |
| **29** | α-D-Glucose | + | + |
| **30** | D-Mannose | + | + |
| **31** | L-Arginine | - | - |
| **32** | L-Histidine | - | - |
| **33** | D-Saccharic Acid | - | - |
| **34** | L-Lactic Acid | + | + |

### Genome assembly parameters

The genome size of novel isolates, PPL117, PPL118^T^, PPL124, PPL125, and PPL126, is approximately 4.86 Mb assembled into 32 to 37 contigs. The genome coverage ranges from 716x to 1434x with 100% genome completeness and 69.2% G+C content. The N50 values range from 230,034 bps to 294,557 bps. The genomes have 4061 to 4065 coding sequences and 50 to 51 tRNA encoding gene sequences (**Table 2**). The genome size, G+C content, and coding sequences of PPL118^T^ are comparable to *X. sontii* PPL1^T^ (Bansal *et al*., 2021).

**Table 2:** Genome assembly and annotation statistics of *Xanthomonas* isolates from this study.

| Strains | Genome size | G+C content (%) | Genome coverage (x) | Number of contigs | N50 (bp) | Genome completeness/contamination | Coding sequences (CDSs) | rRNA/tRNAs | Accession number |
| --- | --- | --- | --- | --- | --- | --- | --- | --- | --- |
| **PPL118<sup>T</sup>** | 4866170 | 69.18 | 787 | 37 | 260645 | 100/0.55 | 4065 | 3/51 | JAQJCQ000000000 |
| **PPL117** | 4865587 | 69.17 | 742 | 32 | 294557 | 100/0.55 | 4065 | 3/51 | JAQQHT000000000 |
| **PPL124** | 4867171 | 69.17 | 849 | 35 | 230034 | 100/0.55 | 4061 | 3/51 | JAQQHW000000000 |
| **PPL125** | 4865229 | 69.17 | 1434 | 33 | 294557 | 100/0.55 | 4065 | 3/50 | JAQQHX000000000 |
| **PPL126** | 4865728 | 69.17 | 716 | 37 | 263124 | 100/0.55 | 4064 | 3/51 | JAQQHY000000000 |

### Genome based phylogenetic and taxonomic characterization

A total of 303 core genes of 24 genomes were used for a mid-rooted phylogeny construction. As observed in many previous studies, Xanthomonads were forming two major clades, clade I with non-pathogenic species such as *X. indica* and *X. sontii*, and pathogens such *X. translucens* and clade II with intensely studies major pathogens of important crops such as *X. oryzae*, and *X. arboricola* (Koebnik *et al*., 2021, Mafakheri *et al*., 2022). The novel isolates, PPL117, PPL118^T^, PPL124, PPL125, and PPL126, are forming a subclade in clade I in close proximity to *X. sontii*, *X. indica*, and *X. sacchari*. *X. sontii* and *X. indica* are also two non-pathogenic *Xanthomonas* (NPX) species reported from healthy rice seeds from India (**Fig. 2**). The novel isolates are forming a monophyletic clade as outliers of *X. sontii* isolates.

**Figure 2.**
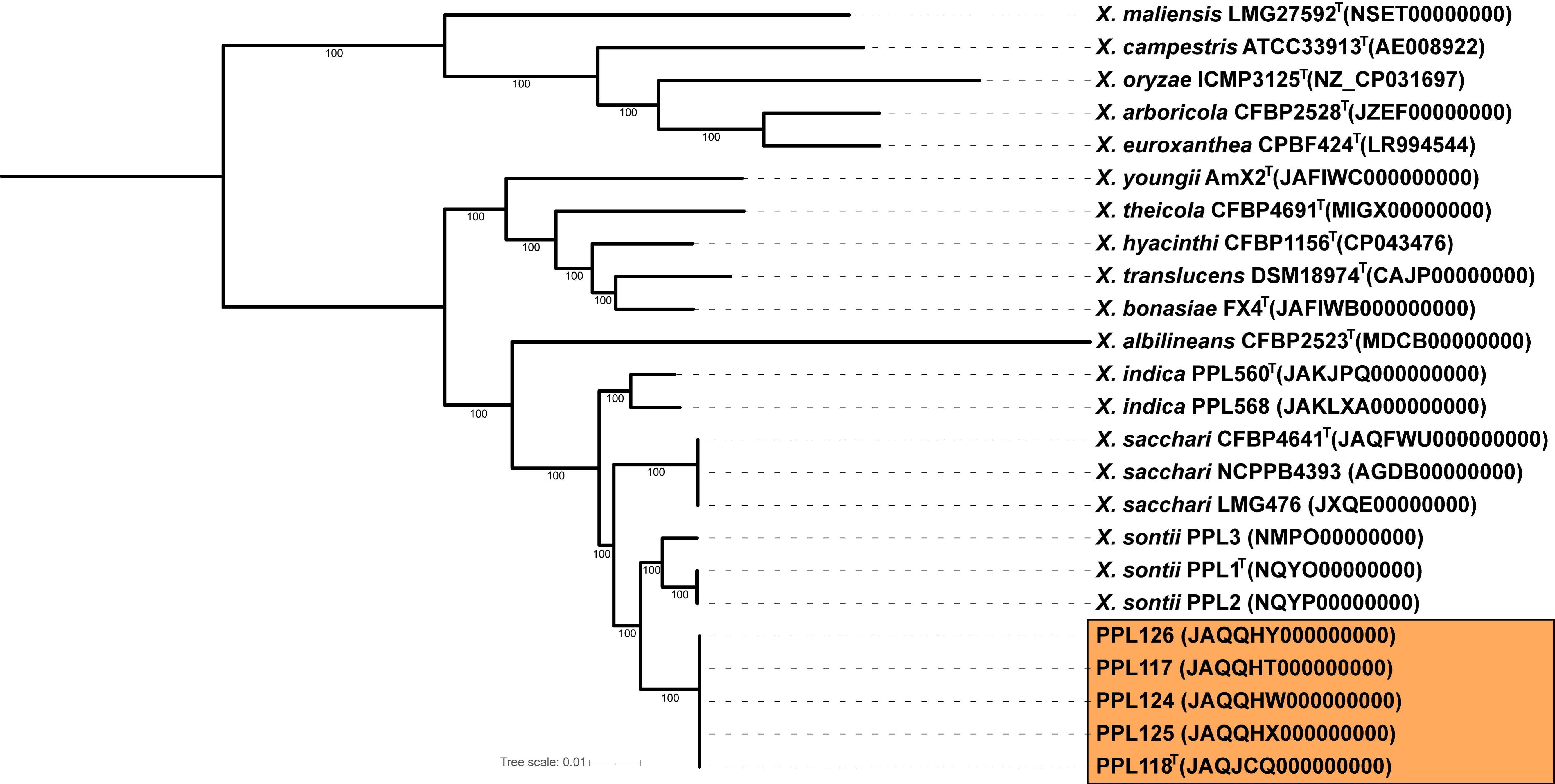
A mid-rooted maximum likelihood phylogeny generated using core genes of the novel *Xanthomonas* isolates with representative of the genus *Xanthomonas*. The bootstrap values above 50% are given at the tree branches. The novel isolates from this study are highlighted in the orange colour. The tree scale represents the number of nucleotide substitutions per site.

Initially, the taxonomic classification of the genus *Xanthomonas* was very complex leading to characterization of up to 100 species which was based on naming new species based on the host (Vauterin *et al*., 2000). With the reduced cost of genome sequencing, the species delineation has become reproducible, fast and accurate. Average Nucleotide Identity (ANI) and digital DNA-DNA hybridization (dDDH) are modern gold standards for species delineation (Chun *et al*., 2018). A combination of cut-off values below 96% for ANI and below 70% dDDH is set for defining a new species (Konstantinidis *et al*., 2006, Auch *et al*., 2010). However, the genomic analysis revealed existence of some species which have ANI values at the borderline of specified cut-off, in such cases dDDH is much more reliable for species delineation (Orata *et al*., 2016, Atanasov *et al*., 2022, Nouioui *et al*., 2023) also in delineation of sub-species (Meier-Kolthoff *et al*., 2014, Meier-Kolthoff & Göker, 2019). This is because dDDH, unlike ANI estimation is not similarity-type index (Palmer *et al*., 2020) and do not involve fragmentation of sequence data (Chun & Rainey, 2014, Hayashi Sant’Anna *et al*., 2019). Moreover, dDDH correlated better with wet-lab DDH than to ANI (Konstantinidis & Tiedje, 2005, Richter & Rosselló-Móra, 2009). Just use of ANI has also led to report of *X. sontii* strains as *X. sacchari* (Rana & Patil, 2023). Hence when studying community of bacteria that form complex, it is important use multiple approaches to establish taxonomic identity.

Therefore, to confirm the species status of the novel isolates ANI values were calculated using two different methods against the type strains of *X. sontii*, *X. indica* and *X. sacchari* which are phylogenetically close relatives of these strains. The orthoANI values of the novel isolates were approximately 93% and 93.5% calculated against *X. indica* and *X. sacchari* type strains. With *X. sontii* type strain the orthoANI values was approximately 95.8%, slightly below the cut-off value i.e., 96%, for species delineations (**Table 3**). ANIb values were slightly less compared to OrthoANI value where the novel isolates shared 95.3% ANI with *X. sontii* type strain. However, the dDDH values of the novel isolates calculated with the novel isolates are 64.4% which is significantly lower than the threshold dDDH values i.e., 70% specified for species delineation. For novel species characterization dDDH is compare much more robust as compared to ANI. Based on the phylogeny, dDDH and ANI values it can be concluded that these novel strains belong to a novel species that is very close relative of *X. sontii*. Here, the ANI and dDDH amongst the novel isolates are 99.9% and 100%, supporting their monophylum in the genome-based tree.

**Table 3:** dDDH and orthoANI/ANIb values of the *Xanthomonas* strains isolated from this study calculated against each other and with the type strains of *X. indica*, *X. sontii*, and *X. sacchari*. Lower triangle in the table represents orthoANI/ANIb values and upper triangle represents dDDH values. Blue-colored and yellow-colored boxes highlight values above cut-offs for dDDH and ANI, respectively.

|  | Strain | 1 | 2 | 3 | 4 | 5 | 6 | 7 | 8 |
| --- | --- | --- | --- | --- | --- | --- | --- | --- | --- |
| 1 | *X. indica* PPL560<sup>T</sup> |  | 51.9 | 51 | 50.7 | 50.7 | 50.7 | 50.7 | 50.7 |
| 2 | *X. sacchari* CFBP4641<sup>T</sup> | 93.6/93.2 |  | 55.1 | 51.6 | 51.7 | 51.7 | 51.6 | 51.7 |
| 3 | *X. sontii* PPL1<sup>T</sup> | 93.3/93.0 | 94.2/93.6 |  | 64.6 | 64.4 | 64.4 | 64.4 | 64.4 |
| 4 | PPL118<sup>T</sup> | 93.3/93.1 | 93.5/93.1 | 95.8/95.3 |  | 100 | 100 | 100 | 100 |
| 5 | PPL117 | 93.2/93.1 | 93.5/93.1 | 95.8/95.3 | 99.9/100 |  | 100 | 100 | 100 |
| 6 | PPL124 | 93.2/93.1 | 93.5/93.1 | 95.7/95.3 | 99.9/100 | 99.9/99.9 |  | 100 | 100 |
| 7 | PPL125 | 93.3/93.1 | 93.5/93.1 | 95.8/95.3 | 99.9/100 | 99.9/100 | 99.9/99.9 |  | 100 |
| 8 | PPL126 | 93.3/93.1 | 93.4/93.1 | 95.8/95.3 | 99.9/100 | 99.9/100 | 99.9/99.9 | 99.9/100 |  |

### Genomic features of the novel isolates

The anvi’o pan-genome analysis revealed that the number of unique genes were 158 in PPL118^T^ and 481, 178, and 259 in type strains of *X. sontii*, *X. sacchari*, and *X. indica*, respectively (**Figure 3a**). The large number of unique genes also support the classification of each of these members into distinct species based on whole genome phylogeny and standard genomic indices. The unique genes of PPL118^T^ were furthered investigated to find out important gene clusters. Unique genes from PIQ37_07510 to PIQ37_07550 were encoding for Lipopolysaccharide gene clusters (LPS). The LPS gene cluster in the novel isolates is different from that of *X. sontii*, *X. indica* and *X. sacchari* (**Figure 3b**). LPS gene cluster is reported to have *etfA* and *metB* at either of its ends. The LPS gene cluster of PPL118^T^ is different at the metB side of the gene cluster where 7 unique genes are present including one transposase encoding gene. The *etfA* side of LPS gene cluster is same as that of *X. sacchari* CFBP4641. Some of the unique genes, from PIQ37_09630 to PIQ37_09650 and from PIQ37_10725 to PIQ37_10750 were encoding for Nonribosomal peptide synthetases (NRPS). This was further confirmed using AntiSMASH analysis where PPL118^T^ was found to carry multiple NRPS regions which mostly constitute a large NRPS gene cluster and could not be assembled into one contig due to shortcomings of draft genomes (**Figure 3c**). The same NRPS is also present in PPL117, PPL124, PPl25, and PPL126. NRPS carrying bacteria are reported to carry antagonistic properties against other bacterial species. Further, the unique genes from PIQ37_06115 to PIQ37_06170 and from PIQ37_10500 to PIQ37_10515 encode for large gene clusters which mostly have hypothetical proteins along with iron transport related genes. The unique gene cluster from PIQ37_019095 to PIQ37_01115 have an tRNA encoding gene upstream and variable G+C content (%) hinting at its acquisition through horizontal gene transfer events. Some of the important unique genes along with their GenBank annotation and G+C content (%) are reported in **Table 4**. These unique genes differentiate the novel isolates from their closest species, *X. sontii*.

**Figure 3.**
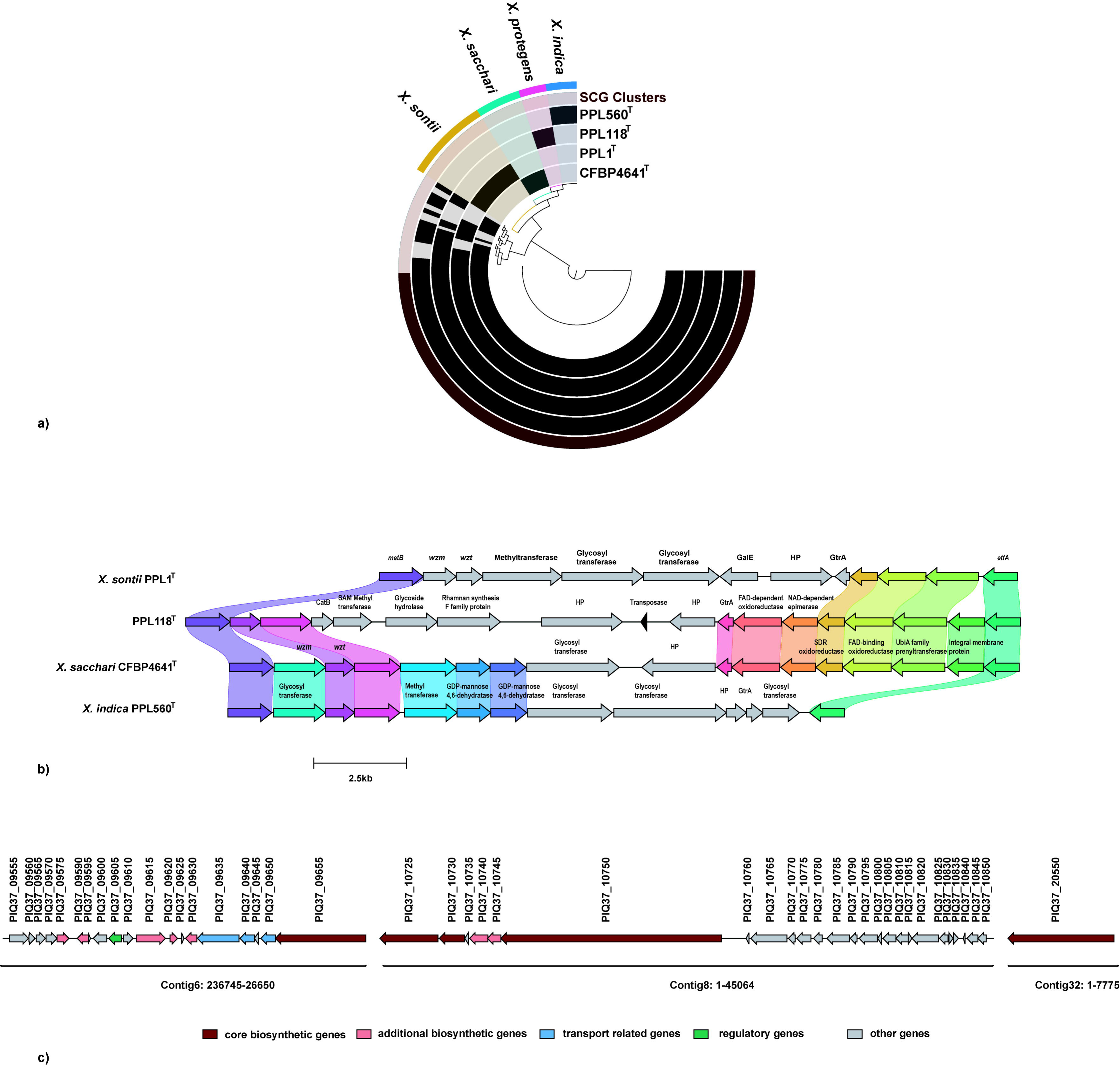
The pan-genome analysis of the PPL118^T^, *X. indica* PPL560^T^, *X. sontii* PPL1^T^ and *X. sacchari* CFBP4641^T^ conducted using Anvio’ platform. **a)** The black colored circular rings represent genomes for each strain. The second outermost ring represents the core genes and the blue, pink, green and yellow colored regions in the outermost ring highlight the unique genes of PPL560^T^, PPL118, CFBP4641^T^, and PPL1^T^. **b)** The Lipopolysaccharide gene cluster comparison of PPL1^T^, PPL118^T^, CFBP4641^T^, and PPL560^T^. The colored arrows depict genes and same gene have same colour across the strains. Unique genes are depicted by grey colored arrows. The colored connecting regions of genes highlight protein homology. **c)** The NRPS encoding genomic region unique to PPL118^T^. The arrows are used to depict the genes and the NCBI locus tags are mentioned on top of arrows. The gene annotations and their genomic coordinates are also mentioned in the figure.

**Table 4:** Unique genes characterizing the outlier species of *Xanthomonas sontii*. The NCBI GenBank locus tag, annotation and GC content (%) of each unique gene is mentioned in the table.

| S. No. | NCBI GenBank Locus tag | GenBank annotation | GC content (%) |
| --- | --- | --- | --- |
| 1. | PIQ37_01095 | type I restriction-modification system subunit M | 61.5 |
| 2. | PIQ37_01100 | restriction endonuclease subunit S | 53 |
| 3. | PIQ37_01105 | DUF262 domain-containing protein | 60.6 |
| 4. | PIQ37_01110 | type I restriction endonuclease subunit R | 59.9 |
| 5. | PIQ37_01115 | SprT family zinc-dependent metalloprotease | 60 |
| 6. | PIQ37_06115 | hypothetical protein | 56.6 |
| 7. | PIQ37_06120 | hypothetical protein | 57.8 |
| 8. | PIQ37_06125 | hypothetical protein | 60.5 |
| 9. | PIQ37_06130 | HEPN-associated N-terminal domain-containing protein | 62.6 |
| 10. | PIQ37_06135 | RES family NAD<sup>+</sup> phosphorylase | 61 |
| 11. | PIQ37_06140 | DUF4238 domain-containing protein | 55.7 |
| 12. | PIQ37_06145 | hypothetical protein | 58.2 |
| 13. | PIQ37_06150 | hypothetical protein | 59 |
| 14. | PIQ37_06155 | hypothetical protein | 58.6 |
| 15. | PIQ37_06165 | FTR1 family protein | 67 |
| 16. | PIQ37_06170 | hypothetical protein | 58 |
| 17. | PIQ37_07510 | ABC transporter permease | 61.8 |
| 18. | PIQ37_07515 | ABC transporter ATP-binding protein | 60.5 |
| 19. | PIQ37_07520 | CatB-related O-acetyltransferase | 64.2 |
| 20. | PIQ37_07525 | class I SAM-dependent methyltransferase | 60.6 |
| 21. | PIQ37_07530 | glycoside hydrolase family 99-like domain-containing protein | 59.8 |
| 22. | PIQ37_07535 | rhamnan synthesis F family protein | 60.5 |
| 23. | PIQ37_07540 | hypothetical protein | 56.8 |
| 24. | PIQ37_07550 | hypothetical protein | 54.4 |
| 25. | PIQ37_09630 | 4'-phosphopantetheinyl transferase superfamily protein | 50.7 |
| 26. | PIQ37_09635 | efflux RND transporter permease subunit | 57.7 |
| 27. | PIQ37_09640 | efflux RND transporter periplasmic adaptor subunit | 60 |
| 28. | PIQ37_09645 | hypothetical protein | 55 |
| 29. | PIQ37_09650 | ABC transporter permease | 62.3 |
| 30. | PIQ37_10500 | alkaline phosphatase D family protein | 68 |
| 31. | PIQ37_10505 | TonB-dependent receptor | 65.2 |
| 32. | PIQ37_10510 | FecR domain-containing protein | 72 |
| 33. | PIQ37_10515 | RNA polymerase sigma factor | 68.2 |
| 34. | PIQ37_10725 | amino acid adenylation domain-containing protein | 66.6 |
| 35. | PIQ37_10730 | ACP S-malonyltransferase | 65.8 |
| 36. | PIQ37_10735 | hypothetical protein | 60.1 |
| 37. | PIQ37_10740 | aminotransferase class I/II-fold pyridoxal phosphate enzyme | 60.9 |
| 38. | PIQ37_10745 | NAD-dependent epimerase/dehydratase family protein | 63.9 |
| 39. | PIQ37_10750 | SDR family NAD(P)-dependent oxidoreductase | 65.5 |
| 40. | PIQ37_17585 | S8 family serine peptidase | 62.9 |
| 41. | PIQ37_17590 | hypothetical protein (adhesin) | 66.3 |
| 42. | PIQ37_17595 | ATP-binding protein | 59.9 |
| 43. | PIQ37_17600 | response regulator transcription factor | 58.8 |

Further, the Macsyfinder analysis of PPL118^T^ revealed the presence of Type I secretion system, Type II secretion system, Type IV secretion system and type V secretion system. Type III secretion system, its effectors and Type VI secretion system which are known to be important pathogenicity factors in the genus *Xanthomonas* were absent in PPL118^T^. The absence of major pathogenicity regions hinted that PPL118^T^ follows non-pathogenic lifestyle same as its relatives, i.e., *X. sontii* and *X. indica*.

### PPL118^T^ is non-pathogenic in nature and provides protection against bacterial leaf blight

Since PPL118^T^ is an outlier of Non-pathogenic species, *X. sontii* and lacks T3SS, we assumed it to be non-pathogenic. Their non-pathogenicity status was confirmed against the host of isolation by leaf clip inoculation. As expected after 14 days of inoculation no disease lesion was observed in the rice leaves compared to the pathogenic strain, *X. oryzae* BXO43 **(Figure 4A)**. Further, PPL118^T^ was also co-inoculated with the pathogen to the rice leaves to check if it inhibits disease symptoms. It was observed after 14 days that no disease lesion was developed in the leaves co-inoculated with both the pathogen and PPL118^T^ **(Figure 4B)**. The disease inhibition property is similar to as reported in its phylogenetically close relatives *X. indica* and *X. sontii*, both of which are also part of healthy rice microbiome (Rana *et al*., 2023).

**Figure 4.**
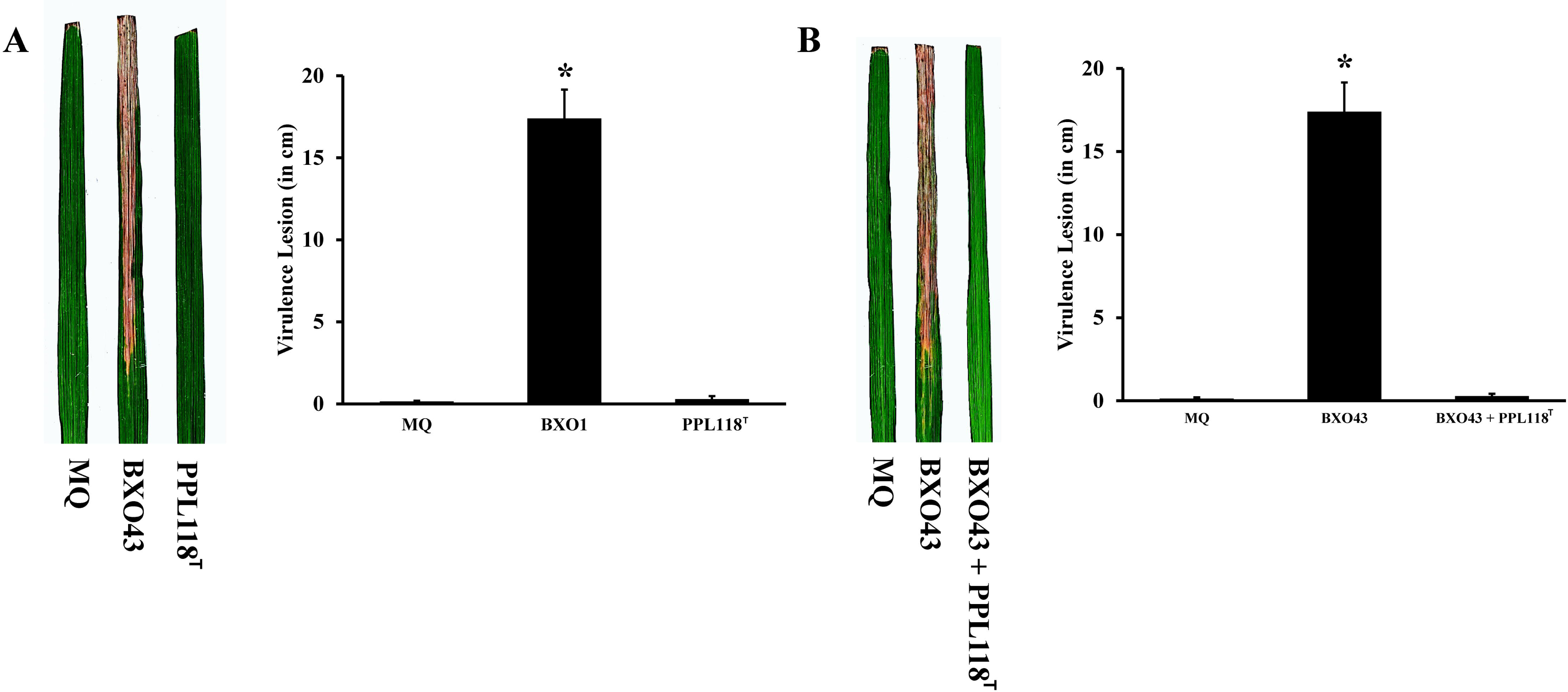
The plant inoculation experiments of PPL118^T^ for **A)** virulence against rice plants **B)** inhibition of disease symptoms due to *X. oryzae*. The Left-hand sides of both the images shows 14 days old rice leaves inoculated with **A)** PPL118^T^ **B)** PPL118^T^ co-inoculated with BXO43, MQ as negative control and BXO43 as negative control. The right-hand sides of the images show disease lesions in cm measured 14 days after inoculation. The error bars depict standard deviation and * labelled column indicates significance difference in lesion length using unpaired two tailed student’s t-test (*p* value < 0.001).

## Conclusion

Our study reinforces the emerging hypothesis that NPX are important group of associated with diverse plants. This is considering the importance of NPX in bio-protection and also probiotic properties. Hence it is important to isolate and characterize all the isolate and characterize all the NPX members of rice plant in general and rice seeds in particular, as the seed microbes are the first in habitants and are associated with the plant throughout their lifecycle. Our study suggests NPX in rice seeds form a species complex and they protect rice plants against pathogenic species. This observation is also supported by genomic investigation as revealed by absence of T3SS/effectome and also presence of a NRPS in the new species. Further using phylogenomics, pangenome and taxono-genomic indices are required when reporting new species and also studying new strains of NPX from a host plant. As the new species is taxonomic outlier of X. sontii, further comparative genomic and molecular studies will help in understanding the success of X. sontii as a keystone species of rice microbiome.

### Description of Xanthomonas gujaratensis sp. nov

*Xanthomonas* protegens (pro.te’gens. L. part. adj. *protegens*, protecting, as the bacterium protects rice plants against the pathogen *Xanthomonas oryzae* pv. *oryzae*)

The strains of this new species appear as pale-yellow colored, mucoid, convex colonies on Nutrient Agar media after 24h of growth at 28 °C. The cells are rod shaped with single polar flagella. The species belongs to the genus *Xanthomonas* which are known to be gram negative. The type strain shows positive growth on pH6, 1% NaCl, 1% sodium lactate, and negative growth on pH5, 8% NaCl. It shows metabolism of Dextrin, D-Maltose, D-Trehalose, D-Cellobiose, Gentiobiose, Sucrose, D-Turanose, α-D-lactose, D-melibiose, β-methyl-D-glucoside, D-Salicin, N-Acetyl-Glucosamine, α-D-glucose, D-mannose, D-fructose, D-galactose, L-Fucose, Glycerol, Gelatin, L-alanine, L-aspartic acid, L-glutamic acid, Pectin, Quinic acid, Methyl pyruvate, L-lactic acid, Citric acid, L-Malic acid, Bromo-succinic acid, Propionic acid, Tween 40. The type strain is not pathogenic to rice plants and inhibits disease lesion development of *X. oryzae*, a rice pathogen upon co-inoculation in rice leaves. The approximate genome size and GC content of the novel isolates is 4.9Mb and 69%, approximately.

The type strain is PPL118^T^=MTCC 13396=CFBP 9164=ICMP 25181.

## Data availability

The partial 16S rRNA gene sequences of PPL117, PPL118^T^, PPL124, PPL125, and PPL126, are submitted to NCBI GenBank database with the accession numbers respectively PP406515, PP406533, PP406535, PP406536, and PP406471, respectively. The whole genome sequences of PPL117, PPL118^T^, PPL124, PPL125, and PPL126, are submitted to NCBI GenBank under the accession numbers JAQQHT000000000, JAQJCQ000000000, JAQQHW000000000, JAQQHX000000000, JAQQHY000000000, respectively. The type strain PPL118 pure culture is submitted in MTCC, CFBP, and ICMP with the accession numbers, PPL118^T^ = MTCC 13396 = CFBP 9164 = ICMP 25181

## Acknowledgements

We acknowledge the funding from a project titled “Deciphering the mechanism(s) of host- endophytes co-evolution, enhanced secondary metabolite production and crop productivity”, grant Number: NBRI-IMTECH-MLP48 and another project entitled OLP-191 titled “Community genomic based insights into adaptation of Xanthomonas to rice and its micro-habitat”. We acknowledge the Council of Scientific and Industrial Research (CSIR) fellowship to RR. We acknowledge Dr Anu Singh for bringing rice seeds from Gujarat.

## Author contributions

RR did the strain isolation, identification, genome assembly, genomic analysis, and drafted the manuscript. AS did phenotypic characterisation and helped with strain isolation. VNM performed plant experiments. RVS and HKP planned and participated in designing of the study. PBP conceived, planned, and participated in designing of study and finalizing the manuscript.

## Conflict of interest

The authors declare no conflict of interest.

---

**Supplementary Figure 1.** Transmission Electron Microscopy (TEM) image of PPL118^T^ (Scale bar = 0.5 µm)

**Supplementary Figure 2.** Biochemical profile of PPL118^T^ calculatedusing BIOLOG GEN III microplate having various tests for utilization of different carbon sources, resistance to different antibiotics, growth at different pH and NaCl concentrations. “+” for positive, “-” for negative and “v” for variable utilization of their respective sources.

## Notes

### Competing Interest Statement

The authors have declared no competing interest.

## References

Abby SS & Rocha EP (2017) Identification of protein secretion systems in bacterial genomes using MacSyFinder. Bacterial Protein Secretion Systems: Methods and Protocols 1–21.

Ahumada GD, Gómez-Álvarez EM, Dell’Acqua M, Bertani I, Venturi V, Perata P & Pucciariello C (2022) Bacterial endophytes contribute to rice seedling establishment under submergence. Frontiers in Plant Science 13: 908349.

Atanasov KE, Miñana Galbis D, Gallego J, Serpico A, Bosch M, Altabella T & Ferrer A (2022) Pseudomonas germanica sp. nov., isolated from Iris germanica rhizomes [Dataset].

Auch AF, von Jan M, Klenk H-P & Göker M (2010) Digital DNA-DNA hybridization for microbial species delineation by means of genome-to-genome sequence comparison. Standards in genomic sciences 2: 117–134.

Bansal K, Kaur A, Midha S, Kumar S, Korpole S & Patil PB (2021) Xanthomonas sontii sp. nov., a non-pathogenic bacterium isolated from healthy basmati rice (Oryza sativa) seeds from India. Antonie van Leeuwenhoek 114: 1935–1947.

Blin K, Shaw S, Augustijn HE, Reitz ZL, Biermann F, Alanjary M, Fetter A, Terlouw BR, Metcalf WW & Helfrich EJ (2023) antiSMASH 7.0: New and improved predictions for detection, regulation, chemical structures and visualisation. Nucleic acids research gkad344.

Buchfink B, Xie C & Huson DH (2015) Fast and sensitive protein alignment using DIAMOND. Nature methods 12: 59–60.

Cesbron S, Briand M, Essakhi S, Gironde S, Boureau T, Manceau C, Fischer-Le Saux M & Jacques M-A (2015) Comparative genomics of pathogenic and nonpathogenic strains of Xanthomonas arboricola unveil molecular and evolutionary events linked to pathoadaptation. Frontiers in plant science 6: 1126.

Chuang S-C, Dobhal S, Alvarez AM & Arif M (2023) Three new species, Xanthomonas hawaiiensis sp. nov., Stenotrophomonas aracearum sp. nov., and Stenotrophomonas oahuensis sp. nov., isolated from Araceae family. bioRxiv 2023.2009.2017.558166.

Chun J & Rainey FA (2014) Integrating genomics into the taxonomy and systematics of the Bacteria and Archaea. International journal of systematic and evolutionary microbiology 64: 316–324.

Chun J, Oren A, Ventosa A, Christensen H, Arahal DR, da Costa MS, Rooney AP, Yi H, Xu X-W & De Meyer S (2018) Proposed minimal standards for the use of genome data for the taxonomy of prokaryotes. International journal of systematic and evolutionary microbiology 68: 461–466.

Cottyn B, Regalado E, Lanoot B, De Cleene M, Mew T & Swings J (2001) Bacterial populations associated with rice seed in the tropical environment. Phytopathology 91: 282–292.

Edgar RC (2004) MUSCLE: multiple sequence alignment with high accuracy and high throughput. Nucleic acids research 32: 1792–1797.

Edgar RC (2010) Search and clustering orders of magnitude faster than BLAST. Bioinformatics 26: 2460–2461.

Eren AM, Esen ÖC, Quince C, Vineis JH, Morrison HG, Sogin ML & Delmont TO (2015) Anvi’o: an advanced analysis and visualization platform for ‘omics data. PeerJ 3: e1319.

Eyre AW, Wang M, Oh Y & Dean RA (2019) Identification and characterization of the core rice seed microbiome. Phytobiomes Journal 3: 148–157.

Fang Y, Lin H, Wu L, Ren D, Ye W, Dong G, Zhu L & Guo L (2015) Genome sequence of Xanthomonas sacchari R1, a biocontrol bacterium isolated from the rice seed. Journal of biotechnology 206: 77–78.

Gilchrist CL & Chooi Y-H (2021) Clinker & clustermap. js: Automatic generation of gene cluster comparison figures. Bioinformatics 37: 2473–2475.

Guindon S, Dufayard J-F, Lefort V, Anisimova M, Hordijk W & Gascuel O (2010) New algorithms and methods to estimate maximum-likelihood phylogenies: assessing the performance of PhyML 3.0. Systematic biology 59: 307–321.

Gurevich A, Saveliev V, Vyahhi N & Tesler G (2013) QUAST: quality assessment tool for genome assemblies. Bioinformatics 29: 1072–1075.

Hayashi Sant’Anna F, Bach E, Porto RZ, Guella F, Hayashi Sant’Anna E & Passaglia LM (2019) Genomic metrics made easy: what to do and where to go in the new era of bacterial taxonomy. Critical reviews in microbiology 45: 182–200.

Koebnik R, Burokiene D, Bragard C, Chang C, Saux MF-L, Kölliker R, Lang JM, Leach JE, Luna EK & Portier P (2021) The complete genome sequence of Xanthomonas theicola, the causal agent of canker on tea plants, reveals novel secretion systems in clade-1 xanthomonads. Phytopathology® 111: 611–616.

Konstantinidis KT & Tiedje JM (2005) Genomic insights that advance the species definition for prokaryotes. Proceedings of the National Academy of Sciences 102: 2567–2572.

Konstantinidis KT, Ramette A & Tiedje JM (2006) The bacterial species definition in the genomic era. Philosophical Transactions of the Royal Society B: Biological Sciences 361: 1929–1940.

Krueger F (2015) Trim Galore!: A wrapper around Cutadapt and FastQC to consistently apply adapter and quality trimming to FastQ files, with extra functionality for RRBS data. Babraham Institute.

Kumar S, Stecher G, Li M, Knyaz C & Tamura K (2018) MEGA X: molecular evolutionary genetics analysis across computing platforms. Molecular biology and evolution 35: 1547.

Lee I, Kim YO, Park S-C & Chun J (2016) OrthoANI: an improved algorithm and software for calculating average nucleotide identity. International journal of systematic and evolutionary microbiology 66: 1100–1103.

Letunic I & Bork P (2021) Interactive Tree Of Life (iTOL) v5: an online tool for phylogenetic tree display and annotation. Nucleic acids research 49: W293–W296.

Mafakheri H, Taghavi SM, Zarei S, Portier P, Dimkić I, Koebnik R, Kuzmanović N & Osdaghi E (2022) Xanthomonas bonasiae sp. nov. and Xanthomonas youngii sp. nov., isolated from crown gall tissues. International Journal of Systematic and Evolutionary Microbiology 72: 005418.

Mafakheri H, Taghavi SM, Zarei S, Rahimi T, Hasannezhad MS, Portier P, Fischer-Le Saux M, Dimkić I, Koebnik R & Kuzmanović N (2022) Phenotypic and molecular-phylogenetic analyses reveal distinct features of crown gall-associated Xanthomonas strains. Microbiology Spectrum 10: e00577–00521.

Martin M (2011) Cutadapt removes adapter sequences from high-throughput sequencing reads. EMBnet journal 17: 10–12.

Meier-Kolthoff JP & Göker M (2019) TYGS is an automated high-throughput platform for state-of-the-art genome-based taxonomy. Nature communications 10: 2182.

Meier-Kolthoff JP, Carbasse JS, Peinado-Olarte RL & Göker M (2021) TYGS and LPSN: A database tandem for fast and reliable genome-based classification and nomenclature of prokaryotes. Nucleic acids research.

Meier-Kolthoff JP, Hahnke RL, Petersen J, Scheuner C, Michael V, Fiebig A, Rohde C, Rohde M, Fartmann B & Goodwin LA (2014) Complete genome sequence of DSM 30083 T, the type strain (U5/41 T) of Escherichia coli, and a proposal for delineating subspecies in microbial taxonomy. Standards in genomic sciences 9: 1–19.

Midha S, Bansal K, Sharma S, Kumar N, Patil PP, Chaudhry V & Patil PB (2016) Genomic resource of rice seed associated bacteria. Frontiers in microbiology 6: 1551.

Niño-Liu DO, Ronald PC & Bogdanove AJ (2006) Xanthomonas oryzae pathovars: model pathogens of a model crop. Molecular plant pathology 7: 303–324.

Nouioui I, Ghodhbane-Gtari F, Pötter G, Klenk H-P & Goodfellow M (2023) Novel species of Frankia, Frankia gtarii sp. nov. and Frankia tisai sp. nov., isolated from a root nodule of Alnus glutinosa. Systematic and Applied Microbiology 46: 126377.

Orata FD, Xu Y, Gladney LM, Rishishwar L, Case RJ, Boucher Y, Jordan IK & Tarr CL (2016) Characterization of clinical and environmental isolates of Vibrio cidicii sp. nov., a close relative of Vibrio navarrensis. International Journal of Systematic and Evolutionary Microbiology 66: 4148–4155.

Page AJ, Cummins CA, Hunt M, Wong VK, Reuter S, Holden MT, Fookes M, Falush D, Keane JA & Parkhill J (2015) Roary: rapid large-scale prokaryote pan genome analysis. Bioinformatics 31: 3691–3693.

Palmer M, Steenkamp ET, Blom J, Hedlund BP & Venter SN (2020) All ANIs are not created equal: implications for prokaryotic species boundaries and integration of ANIs into polyphasic taxonomy. International journal of systematic and evolutionary microbiology 70: 2937–2948.

Parks DH, Imelfort M, Skennerton CT, Hugenholtz P & Tyson GW (2015) CheckM: assessing the quality of microbial genomes recovered from isolates, single cells, and metagenomes. Genome research 25: 1043–1055.

Patil PB & Sonti RV (2004) Variation suggestive of horizontal gene transfer at a lipopolysaccharide (lps) biosynthetic locus in Xanthomonas oryzae pv. oryzae, the bacterial leaf blight pathogen of rice. BMC microbiology 4: 1–14.

Prjibelski A, Antipov D, Meleshko D, Lapidus A & Korobeynikov A (2020) Using SPAdes de novo assembler. Current protocols in bioinformatics 70: e102.

Raj G, Shadab M, Deka S, Das M, Baruah J, Bharali R & Talukdar NC (2019) Seed interior microbiome of rice genotypes indigenous to three agroecosystems of Indo-Burma biodiversity hotspot. BMC genomics 20: 1–16.

Rana R & Patil PB (2023) Xanthomonas sontii, and not Xanthomonas sacchari, is the dominant, vertically transmitted core rice seed endophyte. bioRxiv 2023.2010. 2019.562881.

Rana R, Madhavan VN, Sonti RV, Patel HK & Patil PB (2023) Comparative genomics-based insights into diversification and bio-protection function of Xanthomonas indica, a non-pathogenic species of rice. bioRxiv 2023.2004. 2015.537001.

Rana R, Madhavan VN, Saroha T, Bansal K, Kaur A, Sonti RV, Patel HK & Patil PB (2022) Xanthomonas indica sp. nov., a novel member of non-pathogenic Xanthomonas community from healthy rice seeds. Current Microbiology 79: 304.

Richter M & Rosselló-Móra R (2009) Shifting the genomic gold standard for the prokaryotic species definition. Proceedings of the National Academy of Sciences 106: 19126–19131.

Richter M, Rosselló-Móra R, Oliver Glöckner F & Peplies J (2016) JSpeciesWS: a web server for prokaryotic species circumscription based on pairwise genome comparison. Bioinformatics 32: 929–931.

Ryan RP, Vorhölter F-J, Potnis N, Jones JB, Van Sluys M-A, Bogdanove AJ & Dow JM (2011) Pathogenomics of Xanthomonas: understanding bacterium–plant interactions. Nature Reviews Microbiology 9: 344–355.

Seemann T (2014) Prokka: rapid prokaryotic genome annotation. Bioinformatics 30: 2068–2069.

Triplett LR, Verdier V, Campillo T, Van Malderghem C, Cleenwerck I, Maes M, Deblais L, Corral R, Koita O & Cottyn B (2015) Characterization of a novel clade of Xanthomonas isolated from rice leaves in Mali and proposal of Xanthomonas maliensis sp. nov. Antonie Van Leeuwenhoek 107: 869–881.

Van Dongen S & Abreu-Goodger C (2012) Using MCL to extract clusters from networks. Bacterial molecular networks: Methods and protocols 281–295.

Vauterin L, Rademaker J & Swings J (2000) Synopsis on the taxonomy of the genus Xanthomonas. Phytopathology 90: 677–682.

Wang X, He S-W, He Q, Ju Z-C, Ma Y-N, Wang Z, Han J-C & Zhang X-X (2023) Early inoculation of an endophyte alters the assembly of bacterial communities across rice plant growth stages. Microbiology Spectrum e04978–04922.

Zhang X, Ma Y-N, Wang X, Liao K, He S, Zhao X, Guo H, Zhao D & Wei H-L (2022) Dynamics of rice microbiomes reveal core vertically transmitted seed endophytes. Microbiome 10: 1–19.

