## Supplementary figures and images for "*Xanthomonas protegens* sp. nov., a rice seed-associated probiotic and taxonomic outlier species of *Xanthomonas sontii*"

### Supplementary Fig. 1

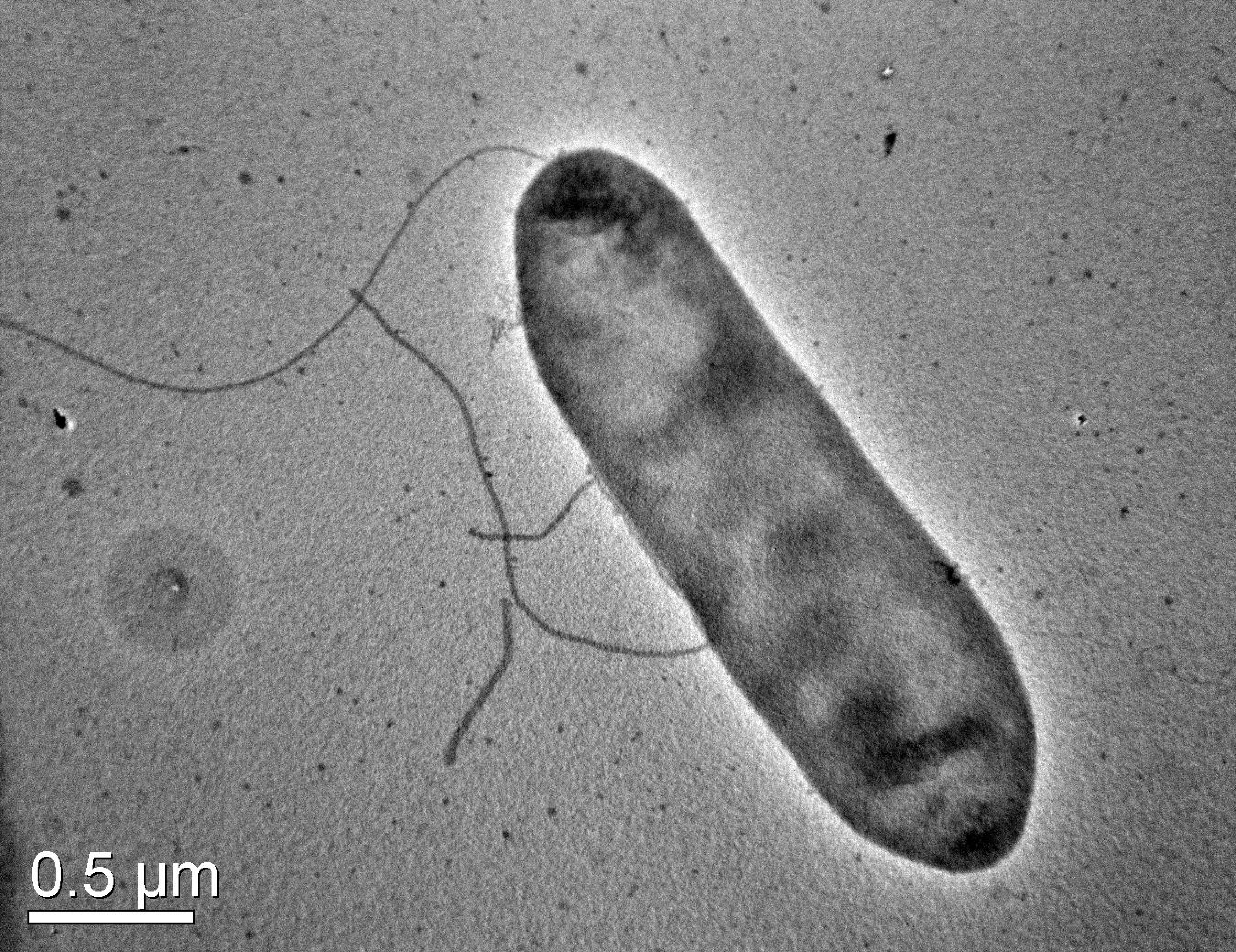

### Supplementary Fig. 2

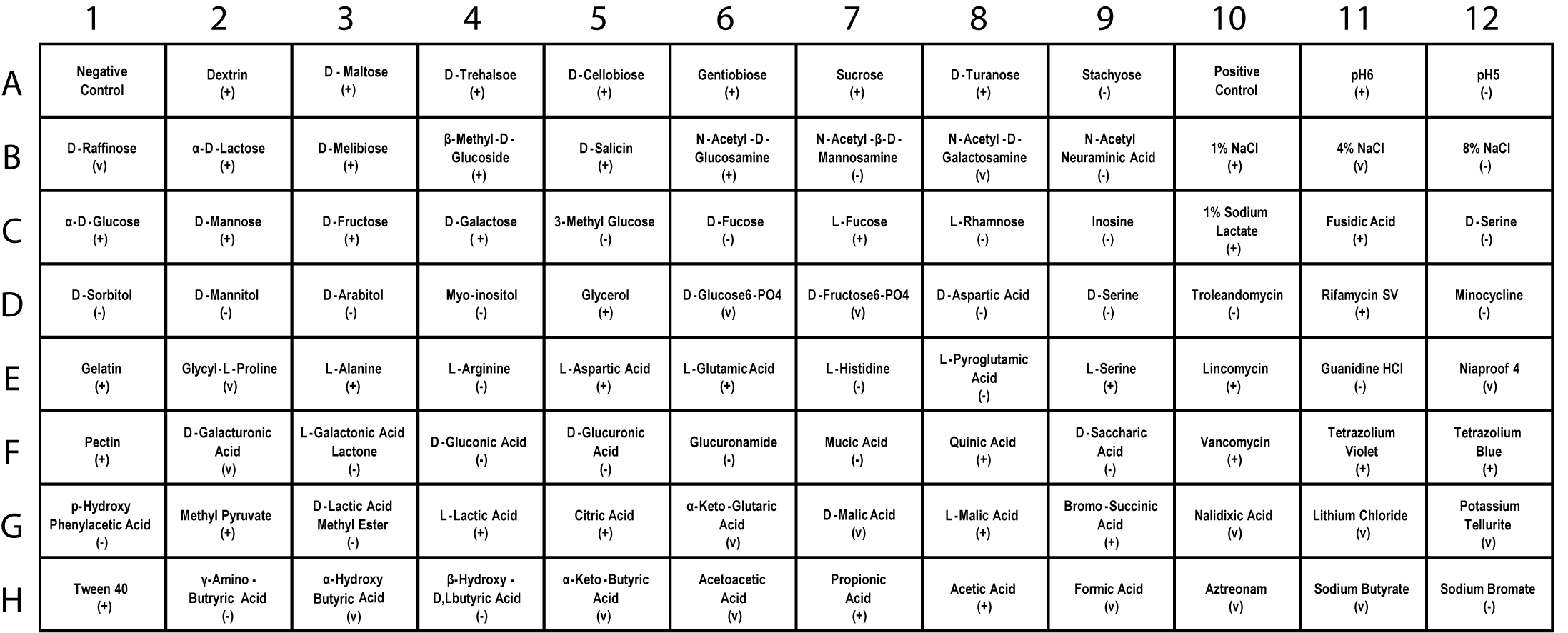
